# Whole-genome sequencing of parvoviruses from wild and domestic animals in Brazil provides new insights into parvovirus distribution and diversity

**DOI:** 10.1101/268219

**Authors:** William Marciel de Souza, Tristan Philip Wesley Dennis, Marcílio Jorge Fumagalli, Jansen de Araujo, Gilberto Sabino-Santos, Felipe Gonçalves Motta Maia, Gustavo Olszanski Acrani, Adriano de Oliveira Torres Carrasco, Marilia Farignoli Romeiro, Sejal Modha, Luiz Carlos Vieira, Tatiana Lopes Ometto, Luzia Helena Queiroz, Edison Luiz Durigon, Márcio Roberto Teixeira Nunes, Luiz Tadeu Moraes Figueiredo, Robert James Gifford

## Abstract

Parvoviruses (family *Parvoviridae*) are small, single-stranded DNA viruses. Many parvoviral pathogens of medical, veterinary and ecological importance have been identified. In this study, we used high-throughput sequencing (HTS) to investigate the diversity of parvoviruses infecting wild and domestic animals in Brazil. We identified 21 parvovirus sequences (including twelve nearly complete genomes and nine partial genomes) in samples derived from rodents, bats, opossums, birds and cattle in Pernambuco, São Paulo, Paraná and Rio Grande do Sul states. These sequences were investigated using phylogenetic and distance-based approaches, and were thereby classified into eight parvovirus species (six of which have not been described previously), representing six distinct genera in the subfamily *Parvovirinae*. Our findings extend the known biogeographic range of previously characterized parvovirus species, and the known host range of three parvovirus genera (*Dependovirus*, *Aveparvovirus*, and *Tetraparvovirus*). Moreover, our investigation provides a window into the ecological dynamics of parvovirus infections in vertebrates, revealing that many parvovirus genera contain well-defined sub-lineages that circulate widely throughout the world within particular taxonomic groups of hosts.

## 1. Introduction

Parvoviruses are small, linear and non-enveloped viruses with single-stranded DNA (ssDNA) genomes ~5-6 kilobases (kb) in length [1]. All parvoviruses possess two major genes, a non-structural (NS) gene encoding the viral replicase, and a capsid (VP) gene encoding the structural proteins of the virion [2]. The *Parvoviridae* family is divided into two subfamilies. All parvoviruses that infect vertebrates falling into one subfamily (*Parvovirinae*), which currently contains 41 viral species, classified into eight genera [1].

Parvoviruses cause disease in humans and domestic animals. For example, parvovirus B19, a species in the genus *Tetraparvovirus*, causes ′erythema infectiosum‵ in children and polyarthropathy syndrome in adults [2], while canine parvovirus, a member of the genus *Protoparvovirus*, can cause haemorrhagic enteritis in dogs, with lethality around 80% of cases [3].

In recent years, high throughput sequencing (HTS) approaches have been instrumental in the discovery of many novel parvovirus species [4–7]. Consequently, the known diversity of parvovirus species has expanded greatly, and recent studies have suggested that parvovirus host range may encompass the entire animal kingdom [8]. To understand the natural biology of vertebrate parvoviruses – i.e. their dynamics in natural hosts, propensity to cause disease, and zoonotic potential – it is important to document their distribution and diversity across a wide range of vertebrate species and populations. In this study, we used a HTS approach to investigate parvovirus infections among wild mammals and birds in Brazil.

## 2. Materials and Methods

### 2.1. Samples

A total of 1073 specimens obtained from 21 different animal species were collected from 2007 to 2016 from rural areas of Pará, Pernambuco, São Paulo, Paraná, Santa Catarina and the Rio Grande do Sul states in Brazil. Individual specimens were distributed in 60 pools based on the species, sample type (i.e., tissue, blood, sera and cloacal swab), date and place of collection (Supplementary Table 1). The species of wild animals were identified using morphological characteristics keys as previously described [9–11]. The geographical distribution of the pools is shown in Figure 1.

### 2.2. Preparation of pools, viral genome sequencing and assembly

Tissues samples were individually homogenized with Hank’s balanced salt solution using the TissueLyser system (Qiagen, USA). Then, the homogenized tissue, sera, and cloacal swabs were centrifuged by 5 min at 10,000g, and the pools were prepared as previously described [12]. The viral genomes were extracted with a QIAamp viral RNA mini kit (Qiagen, USA) and stored at −80°C. Subsequently, the nucleic acid was quantified using a Qubit^®^ 2.0 Fluorometer (Invitrogen, Carlsbad, USA) and the purity and integrity of nucleic acid of samples were measured using an Agilent 2100 Bioanalyzer (Agilent Technologies, USA).

The DNAs were prepared for high-throughput sequencing using the RAPID module with the TruSeq Universal adapter (Illumina, USA) protocols and standard multiplex adaptors. A paired-end, 150-base-read protocol in RAPID module was used for sequencing on an Illumina HiSeq 2500 instrument as recommended by the manufacturer. Sequencing was performed in Life Sciences Core Facility from University of Campinas, Brazil. A total of 7,059,398 to 94,508,748 paired-end reads per pool were generated with 64.85% to 91.45% of bases ≥ Q30 with a base call accuracy of 99.9% (Supplementary Table 1). The sequencing reads were assembled using the de novo approach in the metaViC pipeline (github.com/sejmodha/MetaViC) [12].

**Figure 1.**
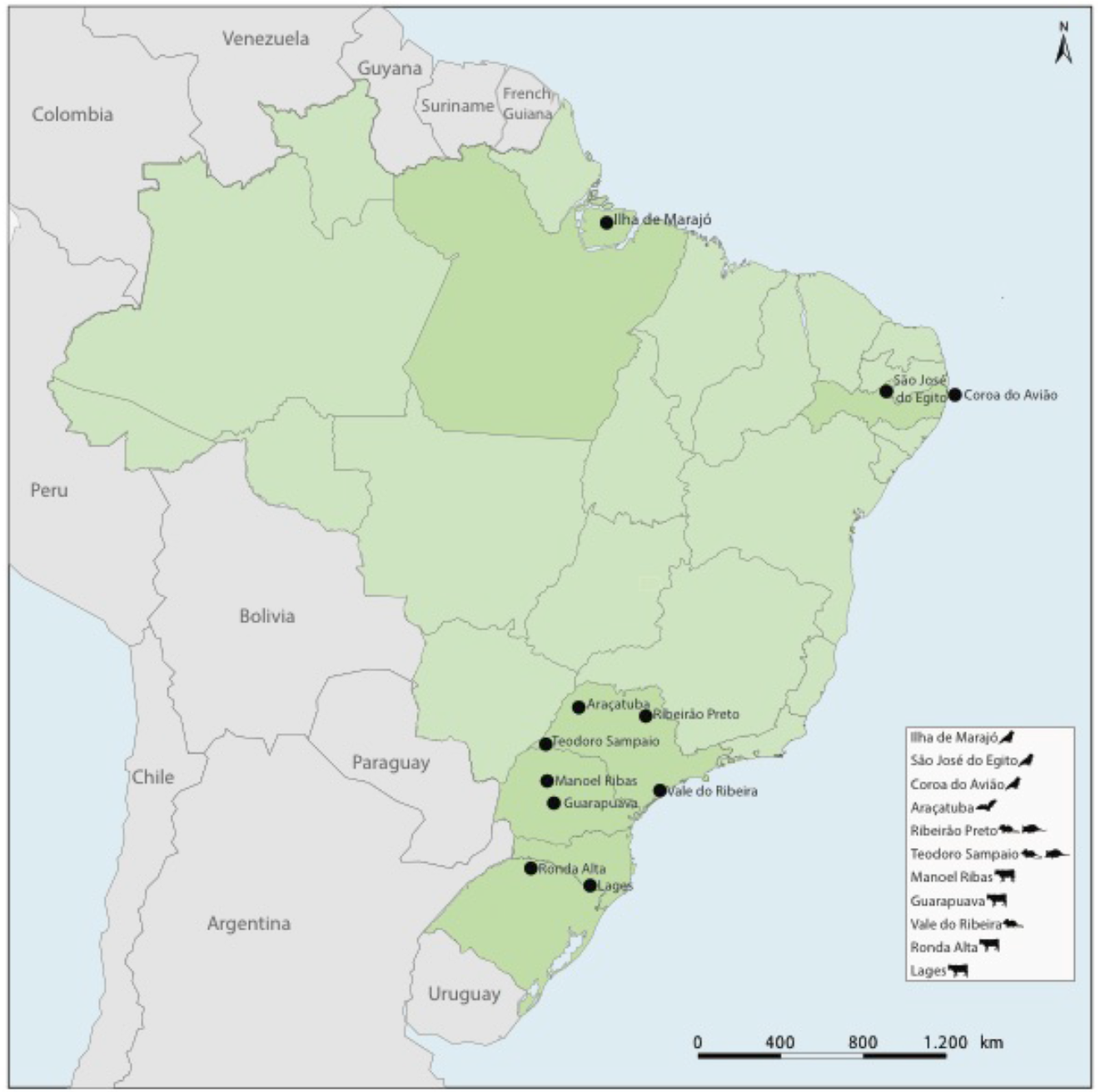
Geographic locations of collected samples in Brazil.

### 2.3. Genome characterization

Genome size, coding potential and molecular protein weight were assessed with Geneious 9.1.2 (Biomatters, New Zealand). The annotations of protein domains were performed using the Conserved Domain Database [13]. The nucleotide sequences determined in this study have been deposited in GenBank under the accession numbers listed in **Table 1**.

### 2.4. Phylogenetic analysis

Maximum likelihood (ML) phylogenetic trees were reconstructed using alignments of NS and VP proteins identified in the present study with representative members of *Parvovirinae* subfamily [1]. Multiple sequence alignment (MSA) was carried out using RevTrans 2.0 [14] with manual adjustment. The alignments of core of NS and VP proteins ML trees were inferred by IQ-TREE version 1.4.3 software based on LG+F+G4 protein substitution model to core of NS protein with 145 amino acids, and LG+F+I+G4 protein substitution model to core of VP protein with 245 amino acids, both with 1,000 replicates [15,16]. Statistical support for individual nodes was estimated via bootstrap replicates. Phylogenetic trees were visualized using Figtree 1.4.2. Nucleotide divergence calculations were performed using the Sequence Demarcation Tool (SDT) version 1.2 in muscle mode [17].

## 3. Results

Using HTS we identified 21 parvovirus sequences in samples derived from rodents, bats, opossums, birds and cattle in Pernambuco, São Paulo, Paraná and Rio Grande do Sul states in Brazil (**Figure 1**). These sequences comprised twelve nearly complete genomes and nine partial genomes (**Table 1**), and included the first examples of parvoviruses identified in opossums, New World bats, and sigmondontine rodents. Parvovirus sequences recovered in our study were classified on the basis of (i) phylogeny and (ii) pairwise distance.

**Table 1.**
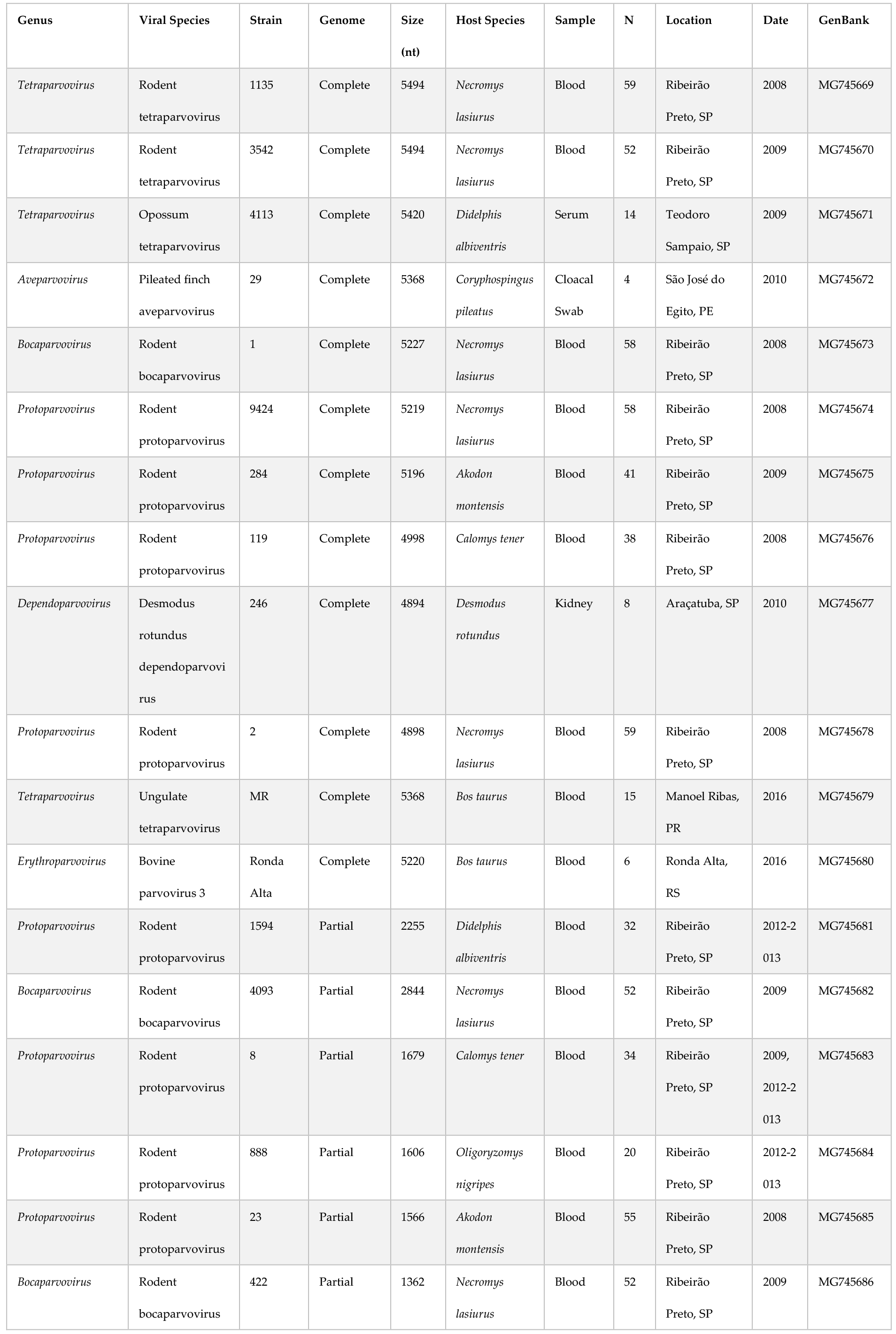
Sequences information, sources, sample, location, location, date and environment of viruses identified in wild animals from Brazil.

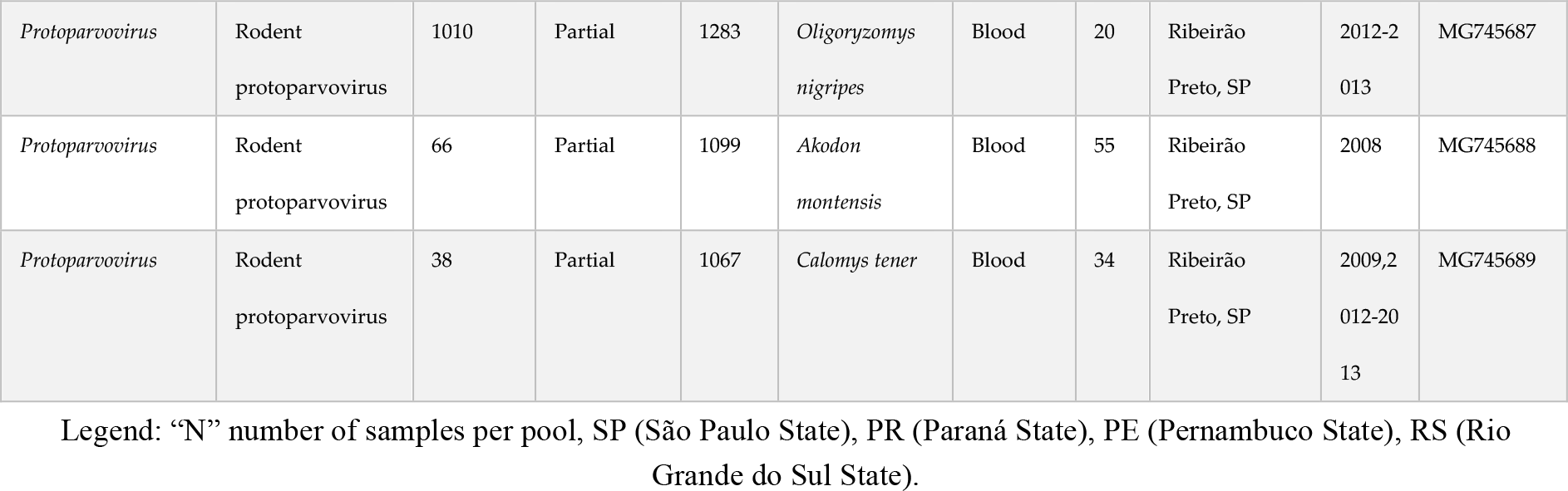

To investigate the phylogenetic relationships of the novel parvoviruses to those described previously, we inferred ML phylogenetic trees from alignments of 71 NS proteins and 71 VP peptide sequences. Phylogenies revealed eight distinct clades corresponding to recognized genera, each having high bootstrap support (values >75%). The sequences recovered in this study grouped into six distinct genera (**Figure 2**). In most cases, the newly identified sequences grouped robustly within the established diversity of their respective genera. Only the *Dependoparvovirus*-like sequence identified in our study grouped in a basal position with respect to previously characterized taxa in both NS and VP trees.

**Figure 2.**
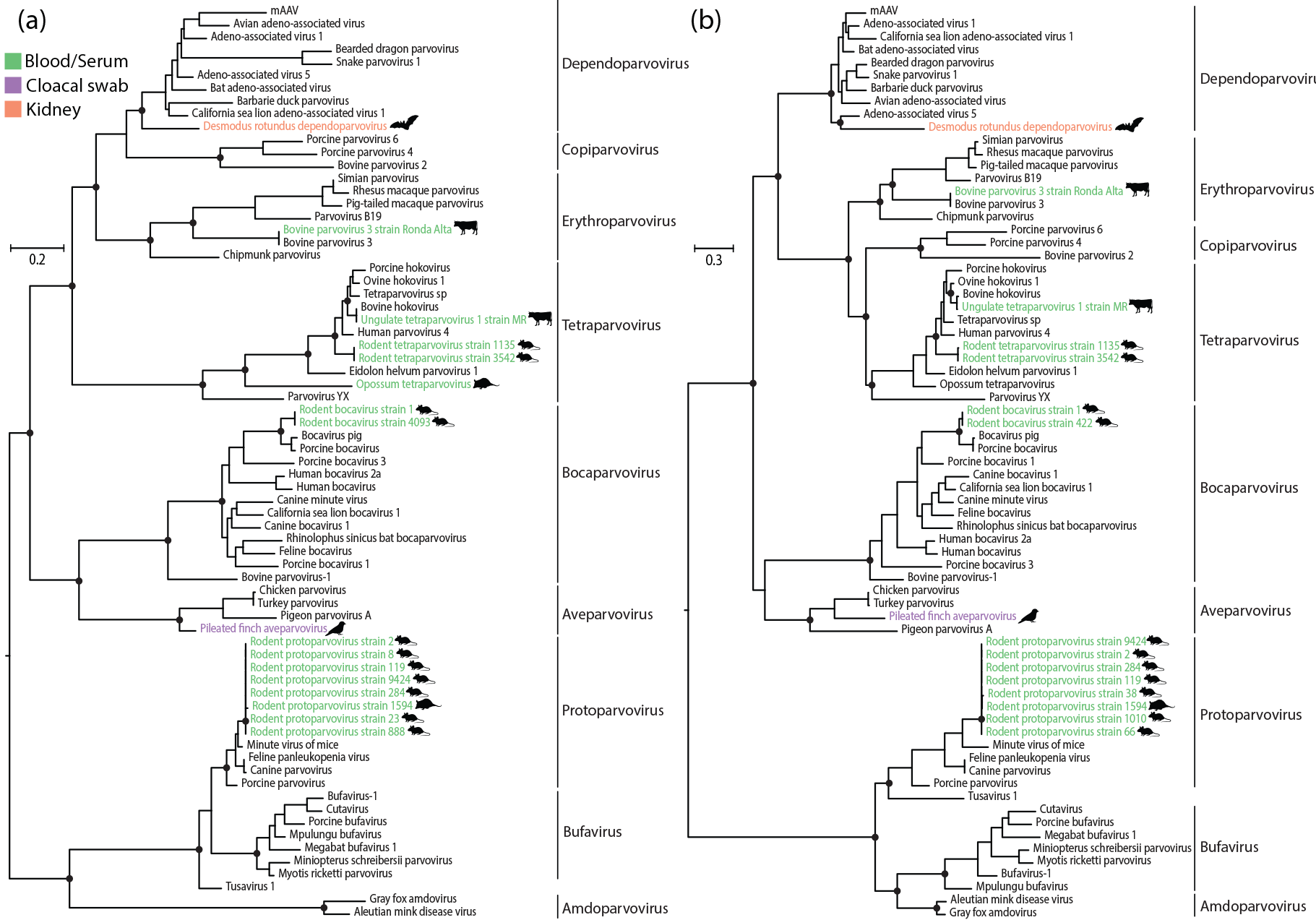
Maximum likelihood phylogenies showing the evolutionary relationships of newly identified parvoviruses. (a) Phylogenetic tree of NS proteins. (b) Phylogenetic tree of VP proteins. Phylogenies are midpoint rooted for clarity of presentation. The scale bar indicates evolutionary distance in substitutions per amino acid site. Black lines indicate genera within the *Parvovirinae* subfamily. Black circles indicate nodes with maximum likelihood bootstrap support levels >75%, based on 1,000 bootstrap replicates. Taxa names of parvoviruses identified in our study are coloured according to sample type, as shown in the key. Silhouettes indicate host species groups.

According to the species demarcation criteria of the International Committee on Taxonomy of Viruses (ICTV), parvoviruses in the same species should share >85% amino acid sequence identity across the entire NS polypeptide sequence [1]. On this basis, the 21 genomes described in this study represent six novel species of parvoviruses, and two that have been described previously (bovine parvovirus 3 and ungulate tetraparvovirus 1) (**Supplementary Figures 1 and 2**).

We identified a novel species of protoparvovirus in sigmondontine rodents. This virus, which was detected in samples from several distinct animal and species (**Table 1**), is quite similar to minute virus of mice (MVM), but is sufficiently distinct based on ICTV criteria to be considered a distinct species. We also identified novel tetraparvoviruses in the opossum and hairy-tailed bolo mouse, and a novel dependoparvovirus in tissue samples derived from common vampire bats (*Desmodus rotundus*). We identified a novel bocaparvovirus species – rodent bocaparvovirus – in two distinct sample pools obtained from hairy-tailed bolo mice, and a novel aveparvovirus in grey pileated finch in São José do Egito, Pernambuco State, Brazil. We also identified strains of two parvoviruses that previously detected in cattle, bovine parvovirus 3 and ungulate tetraparvovirus 1, identified in cattle serum of Ronda Alta in the Rio Grande do Sul State and Manoel Ribas in Paraná State, respectively, both located in South of Brazil.

All the viruses identified in our study have typical parvovirus genome structures encoding NS and VP proteins. The deduced NS protein sequences from these viruses contains the “HxH” and “GPASTGKS” motifs, which play a critical role in viral DNA replication [20]. Most of the capsid proteins also possess the PLA_2_ motif involved in particle release [21], and the glycine-rich (G-rich) region required for cellular entry [22]. Interestingly, we observed that one species, *Desmodus rotundus* dependoparvovirus encodes NS and VP as overlapping ORFs, with a shared region of 47 nucleotides (**Figure 3**).

**Figure 3.**
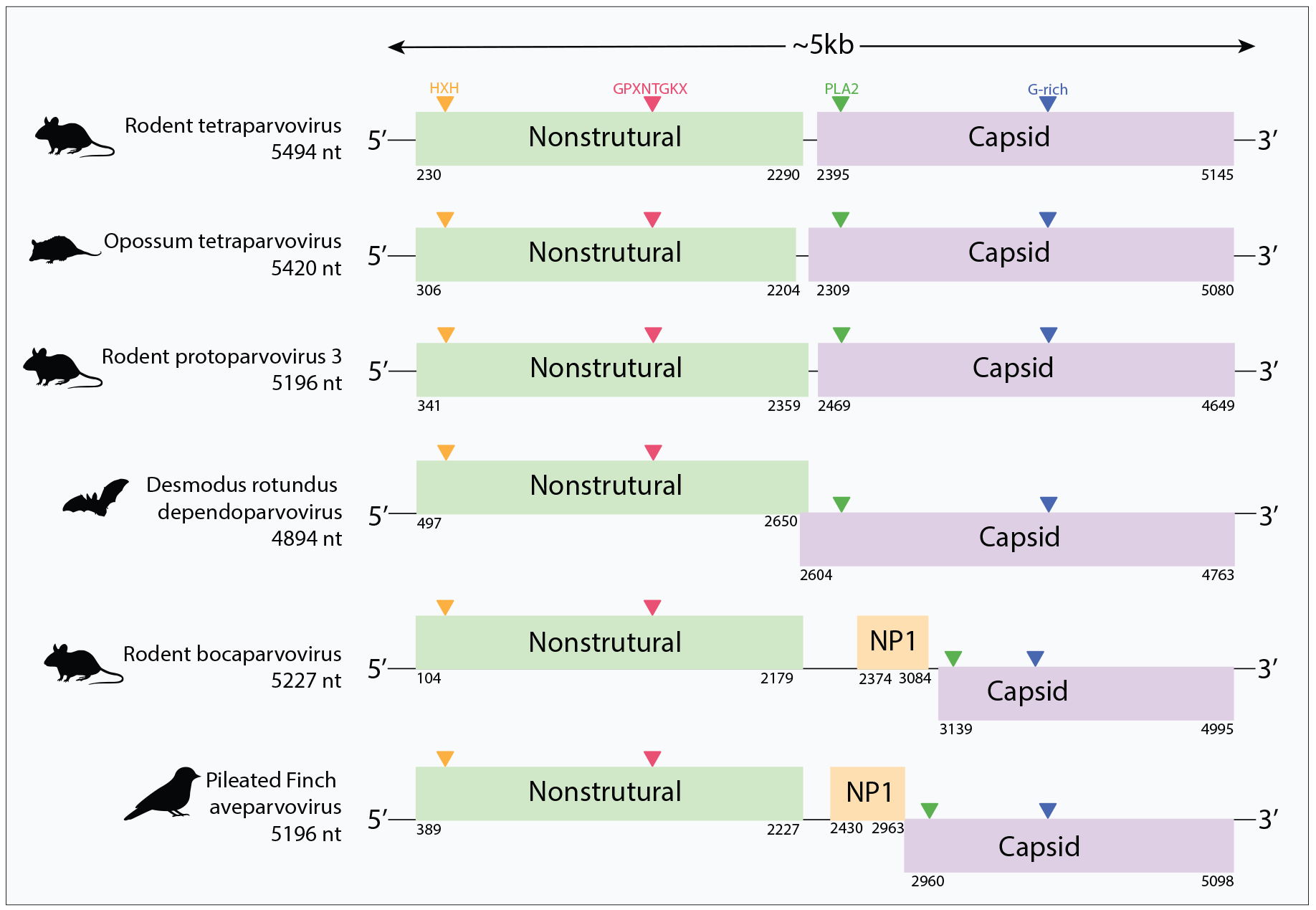
Genome structures of newly identified parvoviruses. The length of the determined nucleotide sequences of the viral sequences are shown in the left. Boxes indicate the open reading frames (ORFs), and the number represent theirs respective position of ORFs.

Notably, the rodent bocaparvoviruses and pileated finch aveparvovirus contain a putative additional ORF (NP1). This gene is located in the middle of the viral genome and overlaps with the C-terminus region of the NS ORF, but in a different reading frame (**Figure 3**). In the case of the rodent bocaparvoviruses, this ORF may correspond to the NP1 protein, which has been reported to play a role in efficient replication for human and canine bocaparvoviruses [25–27], and in immune evasion for porcine bocaparvoviruses [28].

## 4. Discussion

Brazil has a great diversity and abundance of wildlife, and is considered a hotspot for the potential emergence of novel zoonotic viruses [23]. However, parvovirus studies in Brazil have focused predominantly on canine parvovirus and human parvovirus B19 [2,24]. In this study, we used a HTS approach to investigate parvovirus infections among wild mammals and birds apparently without symptoms or disease from Brazil. We identified 21 parvovirus sequences, representing six novel and two previously described parvovirus species. We report the first examples of parvoviruses in samples derived from *Sigmondontinae* rodents, opossums and New World bats. Interestingly, all of the viruses detected here were sequenced from serum or blood samples suggesting that viremia may have been a factor in their identification.

We detected strains of ungulate tetraparvovirus – a virus in the genus *Tetraparvovirus* – in cattle from the South of Brazil. Ungulate tetraparvovirus 2 (formerly known as porcine hokovirus) has previously been identified in swine in Brazil [29]. However, ungulate tetraparvovirus 1 (formerly known as bovine hokovirus) has not previously been reported outside Asia. This virus, which was originally identified in bovine spleen samples obtained from food markets in Hong Kong, has also been identified in domestic yaks (*Bos grunniens*) in northwestern China [18,19]. The identification of this virus in an entirely distinct population (Brazilian cattle) not only establishes that it occurs outside Asia, but also suggests it may be present in cattle populations throughout the world. In addition, we identified novel species of tetraparvovirus in samples obtained from rodents, and from an opossum. Interestingly, the opossum sequence grouped basal relative to the largest *Tetraparvovirus* clade, which contains isolates from diverse eutherian mammals. Further sampling may reveal if this basal position reflects the broad co-divergence of tetraparvoviruses and mammals dating back to the common ancestor of marsupials and eutherians. Such ancient origins of the *Tetraparvovirus* genus are consistent with evidence from endogenous viral element (EVE) sequences that parvoviruses have been infecting mammals for millions of years [30,31].

Recently, studies have reported numerous novel dependoparvoviruses in samples derived from Asian bats [32,33]. Here, we provide the first report of a dependoparvovirus in a New World bat – the vampire bat (*Desmodus rotundus*). In trees based on Rep, this virus groups basally within the *Dependoparvovirus* genus, consistent with these viruses potentially having an ancestral origin in bats, as has been proposed previously [32].

Currently, only one species is recognised in the genus *Aveparvovirus*. This virus (*Galliform aveparvovirus 1*) infects chickens and turkeys and is widespread in poultry farms in the United States and Europe [34,35]. We identified a novel *Aveparvovirus* species in samples derived from pileated finch (*Coryphospingus pileatus*), an indigenous (and non-migratory) South American bird, suggesting that viruses belonging to the *Aveparvovirus* genus may circulate widely among avian species, including wild as well as domestic birds.

We detected bovine parvovirus 3 (genus *Erythroparvovirus*) in Brazilian cattle. Since this virus – to the best of our knowledge – has only been described as a contaminant of commercial bovine serum [36], our study is the first to report detection of bovine parvovirus 3 in cattle populations.

We also identified a novel protoparvovirus species infecting sigmodontine rodents in Brazil. Sigmodontine rodent protoparvovirus was identified in several species of rodents (all belong to subfamily) that we captured in the Ribeirão Preto region of São Paulo State. These viruses are closely related to *Minute virus of mice* (MVM), a common pathogen of laboratory mice [37], but following official taxonomic criteria, they are sufficiently divergent from MVM (>85% in NS and >73% aa in VP) to be considered a distinct species within the *Protoparvovirus* genus.

Bocaparvoviruses are associated with pathogenic conditions in human, bovine and canine hosts [2,38]. Rodent bocaparvoviruses have recently been reported [39], but relatively little is known about their broader distribution. We identified novel rodent bocaparvoviruses in sigmodontine rodents that are closely related to bocaparvoviruses recently reported in brown rats (*Rattus rattus*) in China [39] (data not shown). Together, these findings suggest a broad distribution for rodent bocaparvoviruses.

Parvoviruses that infect domestic and wild carnivores (including amdoviruses and protoparvoviruses) have been studied fairly extensively in the field. These studies have shown that groups of closely related parvoviruses circulate widely among species in the order Carnivora, with the barriers to transmission between species within the order apparently being relatively low [40–42]. The findings of our study suggests that this pattern might be reflected more broadly in parvovirus ecology, with many parvovirus genera containing sublineages that circulate within particular taxonomic groups of hosts (and are largely restricted to this host group). For example, the phylogenetic relationships shown in Figure 1 indicate that closely related protoparvoviruses circulate widely among rodents, and that closely related tetraparvoviruses circulate widely in ungulates. With further sampling of parvovirus diversity it should quickly become apparent whether these inferences are accurate.

## 5. Conclusions

In this study, we used a sequencing-based approach to characterize parvovirus infections in wild and domestic animals in Brazil. Our findings extend the known biogeographic range of previously characterized parvovirus species, and the known host range of three parvovirus genera (*Dependovirus*, *Aveparvovirus*, and *Tetraparvovirus*). More broadly, our findings indicate that many parvovirus genera contain well-defined sub-lineages that circulate widely throughout the world within particular taxonomic groups of hosts.

## Supplementary Materials

: The following are available online at www.mdpi.com/link, Table S1: Samples information, host, sources, sample type, location, date, reads, and %Bases >=Q30. Figure S1: Heatmap of pairwise amino acid identities of the NS protein of parvoviruses identified in this study and representative members of *Parvovirinae* subfamily based on ICTV criteria. The viruses described in this study are highlighted with bold, Figure S1: Heatmap of pairwise amino acid identities of the VP protein of parvoviruses identified in this study and representative members of *Parvovirinae* subfamily based on ICTV criteria. The viruses described in this study are highlighted with bold.

## Acknowledgments

We thank Meire Christina Seki, Janaína Menegazzo Gheller, Luiz Gustavo Betim Góes, Cristiano de Carvalho, Wagner André Pedro, Luciano M. Thomazelli, Fábio Maués, Marcello Schiavo Nardi, Severino M. de Azevedo Júnior, Roberto Rodrigues, Renata Hurtado and Felipe Alves Morais, Márcio Schaefer, Mario Figueiredo, Felipe Morais, Jaqueline R. Silva, Paulo Paulosso, Dercílio Pavanelli, Edison Montilha, Armando F. A. Nascimento, José Teotônio for help in fieldwork. This work was supported by the Fundação de Amparo à Pesquisa do Estado de São Paulo, Brazil (Grant number. 13/14929-1, and Scholarships No. 17/13981-0; 12/24150-9; 15/05778-5; 16/01414-1; 14/20851-8; 06/00572-0; 08/06411-4; 11/06810-9; 11/22663-6; 16/02568-2, 06/00572-0; 09/05994-9 and 11/13821-7). RJG was supported by the Medical Research Council of the United Kingdom (Grant number MC_UU_12014/10).

## Author Contributions

William Marciel de Souza and Robert James Gifford conceived and designed the experiments; William Marciel de Souza, Tristan Philip Wesley Dennis and Marcílio Jorge Fumagalli, Marilia Farignoli Romeiro and Luiz Carlos Vieira performed the experiments; William Marciel de Souza, Tristan Philip Wesley Dennis and Robert James Gifford analyzed the data; Sejal Modha, Márcio Roberto Teixeira Nunes and Luiz Tadeu Mores Figueiredo contributed reagents/materials/analysis tools; Gilberto Sabino-Santos Jr, Felipe Gonçalves Motta Maia, Gustavo Olszanski Acrani, Adriano de Oliveira Torres Carrasco, Luzia Helena Queiroz, Jansen de Araujo, Tatiana Lopes Ometto, Edison Luiz Durigon collected samples and performed fieldwork; William Marciel de Souza, Tristan Philip Wesley Dennis, Luiz Tadeu Moraes Figueiredo and Robert James Gifford wrote the paper.

